# Single point mutations in global regulatory genes restore cephalosporin resistance in a low-MIC *Enterococcus faecium* natural isolate

**DOI:** 10.1101/2025.08.20.671320

**Authors:** Miquel Sánchez-Osuna, Paula Bierge, Inmaculada Gómez-Sánchez, Víctor Monsálvez, Patricia Rabanal, Ana P. Pereira, Ana R. Freitas, Luisa Peixe, Mateu Espasa, Oriol Gasch, Carla Novais, Oscar Q. Pich

**Affiliations:** Laboratori de Recerca en Microbiologia i Malalties Infeccioses, Hospital Universitari Parc Taulí, Institut d’Investigació i Innovació Parc Taulí (I3PT-CERCA), Universitat Autònoma de Barcelona, Sabadell, Spain; Institut de Biotecnologia i Biomedicina, Universitat Autònoma de Barcelona, Bellaterra, Spain; Servei de Malalties Infeccioses, Hospital Universitari Parc Taulí, Institut d’Investigació i Innovació Parc Taulí (I3PT-CERCA), Universitat Autònoma de Barcelona, Sabadell (Spain); UCIBIO, Unidade de Ciências Biomoleculares Aplicadas, Faculdade de Farmácia, Universidade do Porto, Porto, Portugal; Laboratório Associado i4HB - Instituto para a Saúde e Bioeconomia, Faculdade de Farmácia, Universidade do Porto, Porto, Portugal; UCIBIO, Unidade de Ciências Biomoleculares Aplicadas, Instituto Universitário de Ciências da Saúde (1H-TOXRUN, IUCS-CESPU), Gandra, Portugal; Servei de Microbiologia, Hospital Universitari Parc Taulí, Institut d’Investigació i Innovació Parc Taulí (I3PT-CERCA), Universitat Autònoma de Barcelona, Sabadell, Spain

**Keywords:** Ampicillin-susceptible *Enterococcus faecium*, cephalosporin intrinsic resistance, CroS, NusG, RpoB, PBP5

## Abstract

*Enterococcus faecium* exhibits intrinsic resistance to cephalosporins (CPH), yet the genetic determinants of this phenotype remain incompletely understood. To date, *E. faecium* strains with low minimum inhibitory concentrations (MICs) to CPH have only been described following genetic manipulation. At Parc Taulí University Hospital, we identified a clinical isolate of ampicillin-susceptible *E. faecium* (Efm5) that exhibited unusually low-MICs to cefotaxime (1 mg/L), ceftriaxone (3 mg/L), and ceftaroline (0.19 mg/L). Upon single exposure to ceftriaxone (100 mg/L), Efm5 rapidly yielded variants with markedly increased MICs to ceftriaxone (>256 mg/L) and cefotaxime (>32 mg/L), while MICs to ampicillin and ceftaroline were unaffected. Whole-genome sequencing revealed that the high-MIC variants carried single nucleotide polymorphisms (SNPs) leading to non-synonymous mutations in *croS*, *nusG* or *rpoB* genes. Phenotypic assays confirmed that these mutations were associated with ceftriaxone resistance and immunoblots revealed increased expression of penicillin-binding protein 5 (PBP5) in all the high-MIC variants. Transcriptional profiling showed upregulation of the *pbp5* operon, which includes *ftsW*, *psr* and *pbp5*, in the *croS* variants. This study provides the first evidence that *E. faecium* isolates with low-MICs to CPH can arise in clinical settings without laboratory manipulation, challenging the prevailing notion of absolute intrinsic CPH resistance in this species and offering a novel framework to explore previously unrecognized resistance pathways.

## INTRODUCTION

*Enterococcus faecium* is increasingly recognized for its reduced susceptibility to a wide range of commonly used antimicrobial agents, presenting a growing challenge for clinicians in managing infections caused by this pathogen (1). While β-lactams continue to serve as the primary therapeutic agents for susceptible Gram-positive bacteria, *E. faecium* exhibits intrinsic resistance to cephalosporins (CPH), complicating treatment strategies and limiting effective therapeutic options (2). Despite ongoing research, the precise molecular underpinnings of CPH resistance in *E. faecium* remain poorly understood.

CPH resistance in *E. faecium* is largely driven by low-affinity penicillin-binding proteins (PBPs), particularly PBP5 and PBPA. Reduced β-lactam affinity of PBP5 allows cell wall synthesis to continue despite antibiotic exposure (3), while PBPA alters cell wall responses to CPHs (4). Deletion of either protein markedly increases β-lactam susceptibility (3, 4). Conversely, loss of PBPF and PONA, two class-A PBPs, reduces ceftriaxone (CTX) susceptibility while maintaining ampicillin resistance, suggesting a selective role in β-lactam resistance mechanisms (3).

Two-component regulatory systems further contribute to CPH resistance. The CroRS system, well-studied in *Enterococcus faecalis*, regulates genes for cell wall integrity and peptidoglycan turnover, with CPH exposure triggering CroR-dependent *pbp4* overexpression (5–7). The StpA/Stk kinase-phosphatase pair (IreP/IreK in *E. faecalis*) also modulates resistance; mutations enhancing Stk activity or disrupting StpA increase CPH resistance, with StpA loss being particularly impactful in *E. faecium* (8). In *E. faecalis*, *ireK* deletion or combined *ireK/ireP* loss, markedly reduces resistance (9).

To date, *E. faecium* strains with low minimum inhibitory concentrations (MICs) to CPHs have only been obtained through *in vitro* selection, with no reports of naturally occurring isolates. In a recent study, however, we identified several isolates from bacteremic patients with unexpectedly low CPH MICs (10). Here, we investigate one such clinical isolate. Using comparative genomic analysis, we show that non-synonymous mutations in *croS*, *nusG* or *rpoB* restore high-level CPH resistance to the natural isolate with low-MICs, shedding light on previously unrecognized molecular mechanisms that modulate CPH susceptibility in this clinically important pathogen.

## MATERIAL AND METHODS

### Antibiotic susceptibility testing

For research purposes, antibiotic susceptibility tests were conducted using Etest for ampicillin, ceftaroline, CTX, cefotaxime, cefoxitin, vancomycin and rifampicin. Clinical breakpoints for ampicillin, vancomycin and rifampicin were interpreted following the Clinical and Laboratory Standards Institute (11). Clinical breakpoints are not available for CPHs.

### Recovery of CPH high-MIC isolates

One hundred microliters of an Efm5 suspension (10^8^ CFU/mL) from an overnight culture grown without antibiotic selection in Mueller-Hinton (MHB, Merck) medium, were seeded onto Brain Heart Infusion (BHI, Thermo Fisher Scientific) agar plates supplemented with 100 mg/L CTX and incubated at 37°C for 48 hours. Colonies growing in such CTX concentration, from now on called CPH high-MIC isolates, were subsequently selected and subcultured onto plates supplemented with 100 mg/L CTX. Frequency of CPH high-MIC variants was determined by dividing the CFUs counted on BHI agar supplemented with 100 mg/L CTX by the total CFU count detected on BHI agar without antibiotics.

### Whole Genome Sequencing

Whole-genome sequencing of Efm5 and the CPH high-MIC variants was performed using Illumina technology, following the pipeline previously described by our group (10). Multilocus sequence typing (MLST) was performed using the MLST software (https://github.com/tseemann/mlst), based on the recently published MLST scheme for *E. faecium* (12). Putative acquired antibiotic resistance genes were predicted with ABRicate (https://github.com/tseemann/abricate) using the in-built NCBI AMRFinderPlus (13). Prediction of the ampicillin resistance phenotype due to *pbp5* mutations was performed with ResFinder (version 4.7.2) and virulence genes were assessed with Virulence Finder, both from the Center for Genomic Epidemiology (CGE) (14, 15). WGS data are available at National Center for Biotechnology Information (NCBI) under the accession number PRJNA1290041.

### Comparative genomics

To identify potential transposon insertions or premature stop codons responsible for the CPH low-MIC observed in the Efm5 isolate, a BLASTP analysis was performed using the following genes from the *E. faecium* RefSeq reference genome (SRR24) as the queries: *pbp5* (E6A31_RS06825), *pbpA* (E6A31_RS03475), *croR* (E6A31_RS13520), *croS* (E6A31_RS13525), *stpA* (E6A31_RS12560), *stk* (E6A31_RS12555) and *murAA* (E6A31_RS09060), previously reported as key contributors to intrinsic resistance to CPH (4, 8, 16, 17).

On the other hand, sequencing reads of the CPH high-MIC variants were mapped against the Efm5 genome with Snippy (https://github.com/tseemann/snippy). Synonymous Single-Nucleotide Polymorphisms (SNPs) were discarded for further analysis. Predicted mutations were confirmed using Sanger sequencing (Servei de Genòmica i Espectroscòpia de Biomolècules, Universitat Autònoma de Barcelona).

To gain insights into the global distribution of observed mutations across the *Enterococcus* genus, CroS, NusG and RpoB homologs were compiled from the 767 complete *Enterococcus* spp. assemblies available in the NCBI database. This analysis was conducted through reciprocal BLASTP (18) using the *E. faecium* Efm5 CroS, NusG and RpoB protein sequences as the queries, a conservative e-value of <1e–20 and query coverage of >75%, similar to previously described by our group (19). Protein multiple sequence alignments of predicted homologs were performed with CLUSTALW using default parameters (20).

### *In vitro* time-kill curves

Time-Kill Curves (TKCs) were conducted in MHB with (10 mg/L) and without CTX (Sigma Aldrich), using an initial inoculum of 10^6^ CFU/mL. Efm5 and three representative CPH high-MIC variants were incubated at 35°C and 150 rpm. Aliquots of 100 μL were taken at 0, 2, 4, 6 and 24 hours for each strain and condition, and 10 μL of each aliquot was plated in duplicate on BHI agar, which was then incubated at 37°C for 24 hours to determine bacterial viability, expressed as CFUs/mL. Experiments were performed in triplicate following standard procedures and the average of counts was considered (21).

### Immunoblotting

Whole cell lysates were prepared from exponential cultures of *E. faecium* Efm5, the five CPH high-MIC variants and GE-1 (a *pbp5* negative mutant), grown on MHB. Samples were heated at 100°C for 10 min, mixed with Laemmli sample buffer and normalized so that the total protein concentrations were equal. Then, samples were electrophoresed on 10% polyacrylamide gels (SDS-PAGE) by standard procedures. Gels were stained with Coomassie blue or transferred electrophoretically to nitrocellulose membranes following the manufacturer’s protocol. Membranes were then probed with rPBP5-S polyclonal rat sera (1:2,000) followed by HRP-conjugated goat anti-rat IgG antibodies (1:2,000) (RAb-035 Neo Biotech) and developed using Luminata Forte Western HRP substrate (Millipore) following manufacturer’s instructions. In addition to the band corresponding to PBP5 (∼70 kDa), a non-specific lower-molecular-mass band was detected and used as an internal loading control.

### Transcriptional analysis

Exponential cultures of Efm5 and all the CPH high-MIC variants were grown on MHB without antibiotics at 37°C and 150 rpm. Total RNA was purified using the RNeasy Mini Kit (Qiagen) according to the manufacturer’s instructions. At the end, RNA was treated with TURBO DNase (Thermo Fisher) according to the manufacturer’s recommended protocol. Concentration and purity of the extracted RNA were assessed using an RNA Nano Chip (Agilent Technologies).

RT-qPCR was employed to assess gene expression. cDNA synthesis was carried out using the iScriptTM cDNA Synthesis Kit (Bio-Rad) and random primers. qPCR was performed with iTaq polymerase (Bio-Rad) and SYBR green in a CFX96 PCR instrument (Bio-Rad). Relative gene expression was calculated using the Pfaffl method (22). Differential gene expression was judged based on the common arbitrary 2-fold cutoff using *adk*, *gyrB* and *ddl* as housekeeping genes. Data presented in the manuscript corresponds to the analysis of RNAs isolated from three independent biological repeats. Primers used for qPCR were designed using Primer3 software and they are listed in Table S1.

### Electron microscopy

For Scanning Electron Microscopy (SEM) examination, exponential-phase cultures of *E. faecium* Efm5 and three representative CPH high-MIC variants, grown in BHI broth without antibiotics for 2.5 hours at 37°C and 150 rpm, were fixed in a solution containing 2.5% glutaraldehyde in 0.1M phosphate buffer (PB) for 2 hours at 4°C. Then, they were post-fixed in a solution containing 1% osmium tetroxide with 0.8% potassium ferrocyanide for an additional 2 hours. Subsequently, samples underwent dehydration using a series of increasing ethanol concentrations (50%, 70%, 90%, 96% and 100%). After dehydration, chemical drying was performed using hexamethyldisilazane (HMDS) (EMS, Hatfield, PA, USA), followed by sputter coating with a thin layer of PdAu for 4 minutes at 20 mA. Finally, the samples were visualized using a SEM Merlin (Zeiss MErlin, Germany) operated at 5 kV. At least 20 micrographs of each strain were taken and analyzed.

For Transmission Electron Microscopy (TEM) examination, Efm5 and the three representative CPH high-MIC variants were grown, fixed, post-fixed and dehydrated following the same procedure as described for SEM. Next, the cell pellets were embedded in EPON Epoxy Resin (EMS, Hatfield, PA, USA) and polymerized at 60°C for 48 hours. Thin sections with a thickness of 70 nm were obtained using a Reichert-Jung Ultracut E ultramicrotome. These sections were then strained with 2% uranyl acetate and Reynold’s solution (0.2% sodium citrate and 0.2% lead nitrate), followed by examination under a Hitachi H-7000 transmission electron microscope operating at a voltage of 75 kV. All chemicals and reagents were procured from Sigma Chemical Co. (St. Louis, Mo, USA), unless otherwise specified. At least 20 micrographs of each strain were taken and analyzed. Given the particular features observed in the *rpoB*-R5 variant, independent preparations were analyzed.

## RESULTS

### Isolation and clinical context of the *E. faecium* Efm5 strain

An *E. faecium* clinical strain, designated Efm5, was isolated from an 89-year-old female patient with bacteremia at Parc Taulí Hospital in 2016 (10). The patient had a history of cerebrovascular accident with complete dependence, a permanent urinary catheter and a doubtful penicillin allergy. She presented with fever and possible cough; urinary and respiratory tract infections were suspected. Empirical treatment with levofloxacin and clindamycin was initiated. After 24 h, blood cultures were positive in 2 out of 4 bottles: one grew *Escherichia coli* and another yielded *E. coli* plus *E. faecium*. The final diagnosis was healthcare-associated bacteremia, most likely of urinary origin. Based on broth microdilution results, treatment was switched to ceftriaxone (the patient had previously tolerated cefuroxime). Although *E. coli* was susceptible to cephalosporins, no susceptibility testing was performed for *E. faecium*. Since this species is intrinsically resistant to cephalosporins, the treatment decision can be considered inappropriate. Nevertheless, the infection resolved without further complications.

### Genetic characterization of *Enterococcus faecium* Efm5

WGS analysis identified the *E. faecium* Efm5 strain as sequence type 1195 (ST1195) according to PubMLST for *E. faecium*. It was susceptible to vancomycin (MIC = 3 mg/L) and ampicillin (0.25 mg/L), and resistant to rifampicin (MIC = 8 mg/L). It also exhibited low-MICs to ceftaroline (0.19 mg/L), cefotaxime (1 mg/L) and CTX (3 mg/L) (Table 1).

**Table 1:** Minimum Inhibitory Concentrations (MICs, mg/mL) to several antibiotics of the *Enterococcus faecium* Efm5 strain and in representative high-CPH MIC variants (*croS-*R1, *nusG*-R3 and *rpoB*-R5). N/A, not applicable.

| Strains | Ampicillin | Ceftaroline | Ceftriaxone | Cefotaxime | Cefoxitin | Rifampicin | Vancomycin |
| --- | --- | --- | --- | --- | --- | --- | --- |
| Efm5 | 0.25 | 0.19 | 3 | 1 | 48 | 8 | 3 |
| *croS*-R1 | 1 | 0.5 | > 256 | > 32 | 128 | N/A | 4 |
| *nusG*-R3 | 0.38 | 0.19 | > 256 | > 32 | 64 | N/A | 2 |
| *rpoB*-R5 | 1.5 | 0.5 | > 256 | > 32 | 64 | > 32 | 2 |

Several acquired resistance genes were predicted, including those conferring resistance to chloramphenicol (*catA8*), macrolides (*msrC*) and tetracyclines [*tet(L)*, *tet(M*)]. Notably, genes associated with CPH resistance (*pbp5*, *pbpA*, *croRS*, *stpA-stk*, *murAA*) were identified without any transposon insertions or nonsense mutations that could lead to premature protein termination. However, our *in silico* analyses revealed a nonsense mutation in the *psr* gene. Mutations in PBP5 predicted to be associated with ampicillin resistance by ResFinder included the V24A, S27G, R34Q, G66E, E100Q, K144Q, T172A, L177I, A216S, T324A, N496K, A499I and E525D. VirulenceFinder analysis revealed the presence of multiple virulence factor-encoding genes, such as *acm*, *bepA*, *ccpA*, *empABC*, *fms13*, *fms14*, *fms15*, *fms17*, *fnm*, *sagA* and *scm*.

### Isolation and phenotypic characterization of CPH high-MIC variants

Five CPH high-MIC colony variants (R1-R5) of Efm5 were detected in BHI agar plates supplemented with CTX (100 mg/L), yielding an isolation frequency of 5 × 10⁻⁷. Etest results confirmed that they exhibited elevated MICs to CTX (>256 mg/L), cefotaxime (MIC >32 mg/L) and cefoxitin (>64 mg/L) compared to the parental Efm5 strain (Table 1). However, all variants remained susceptible to ampicillin and vancomycin, and with low-MICs to ceftaroline. Notably, the R5 variant also exhibited a 4-fold increase in the MIC to rifampicin (Table 1).

Comparative genomics revealed that each CPH high-MIC variant harbored one SNP compared to the parental Efm5 strain (Figure 1). Specifically, the R1 and R2 variants contained point mutations in the *croS* gene, resulting in V171A (*croS-*R1 variant) and R343H (*croS-*R2 variant) substitutions in the CroS protein (Figure 1A). Structural analysis using AlphaFold predictions indicated that both V171 and R343 are located near the phosphoryl-accepting residue H173 of CroS, which is critical for the activity of this regulatory protein (Figure 1B) (5).

**Figure 1:**
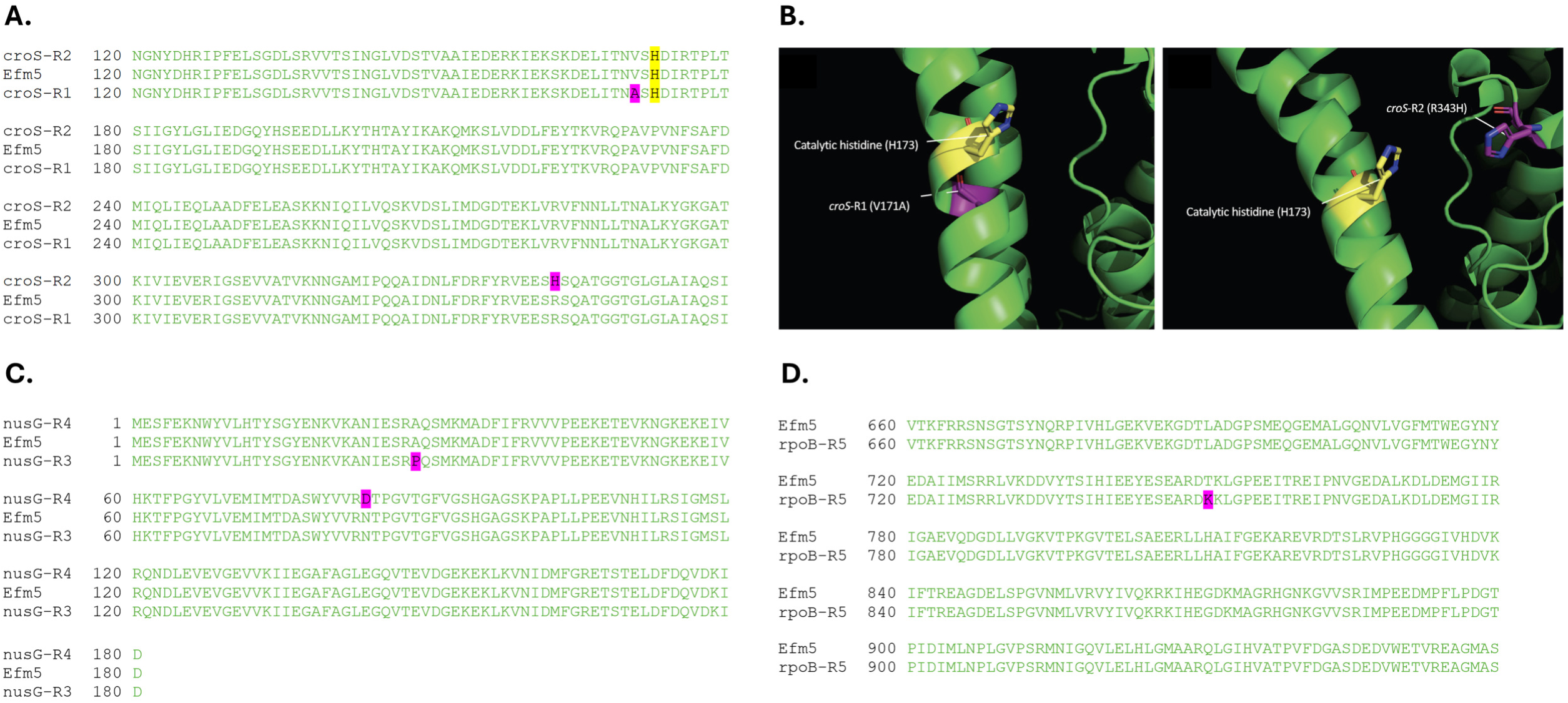
(A) Segment of the CroS multiple sequence alignment encoded in Efm5, *croS*-R1 and *croS*-R2 variants. The catalytic histidine (H173) is highlighted in yellow. (B) Visual representation of the CroS protein 3D structure for the *croS*-R1 (left) and *croS*-R2 (right) variants, generated by AlphaFold (Q3XYJ6) and visualized with PyMOL. (C) NusG multiple sequence alignment encoded in Efm5, *nusG*-R3 and *nusG*-R4 variants. (D) Segment of the RpoB multiple sequence alignment encoded Efm5 and the *rpoB*-R5 variant. In all panels, predicted mutations are labeled in purple.

The R3 and R4 variants presented mutations in the *nusG* gene, leading to A29P (*nusG-*R3 variant) and N84D (*nusG-*R4 variant) substitutions in the NusG protein (Figure 1C). NusG is a conserved intrinsic transcription termination factor essential for transcriptional regulation in bacteria (23). Finally, the rifampicin resistant R5 variant exhibited a T798K substitution in the RpoB protein (Figure 1D), which encodes the β-subunit of RNA polymerase (*rpoB*-R5 variant).

The search for homologs of such variants of CroS, NusG and RpoB in all complete *Enterococcus* genome assemblies revealed that over 99.0% of genomes encoded *croS*, *nusG* and *rpoB* genes. Identical NusG, RpoB and CroS protein sequences to those found in Efm5 strain were identified in 48.2%, 36.1% and 40.4% of the genomes analyzed, respectively. However, NusG, RpoB and CroS sequence variants associated with high-MIC values to CPHs in our isolates were not detected in any of the analyzed genomes in the NCBI database (Table S2). The absence of these mutations in publicly available NCBI genomes supports the notion that the high-MIC variants may harbor compensatory changes offsetting the genetic alterations underlying the unusually low CPH MICs observed in the Efm5 isolate.

### Time-kill curves

Time-kill curves were conducted in the presence of 10 mg/L of CTX. The results confirmed the predicted inhibitory effect of CTX on Efm5 (Figure 2). In contrast, the *croS*-R1, *nusG*-R3 and *rpoB*-R5 variants were all able to grow in the presence of CTX.

**Figure 2:**
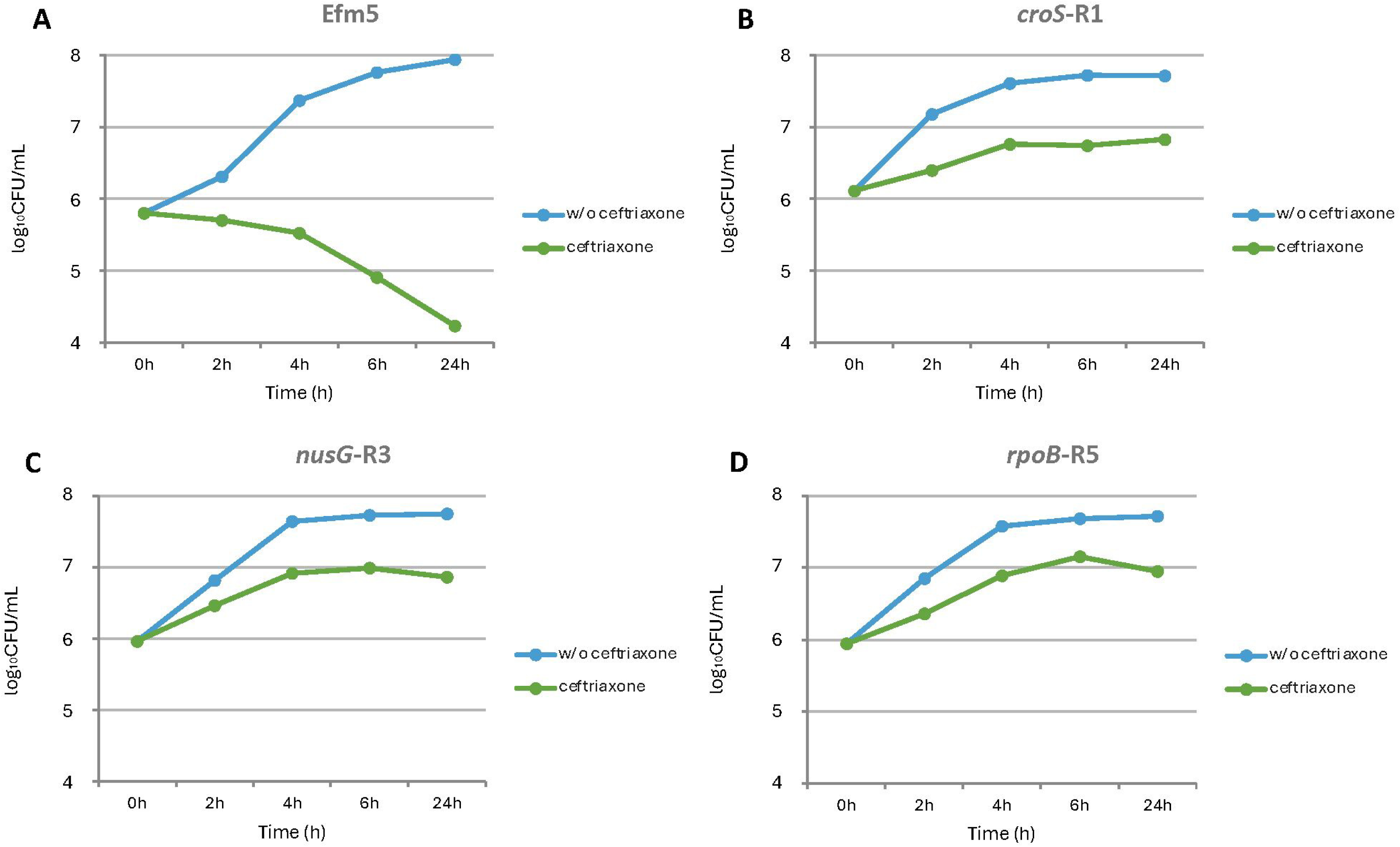
Growth curves of the (A) CPH low-MIC Efm5 and the CPH high-MIC variants (B) *croS*-R1, (C) *nusG*-R3, and (D) *rpoB*-R5, both in absence and presence of ceftriaxone (10 mg/L).

### Assessment of differential expression

As PBP5 is the primary determinant of CPH resistance in *E. faecium* (3), its expression was assessed via Western blotting using an anti-PBP5 monoclonal antibody (24). Our results revealed increased levels of PBP5 expression in all the CPH high-MIC variants compared to the Efm5 strain (Figure 3). No PBP5 was detected in *E. faecium* GE-1, a *pbp5* negative strain (25).

**Figure 3:**
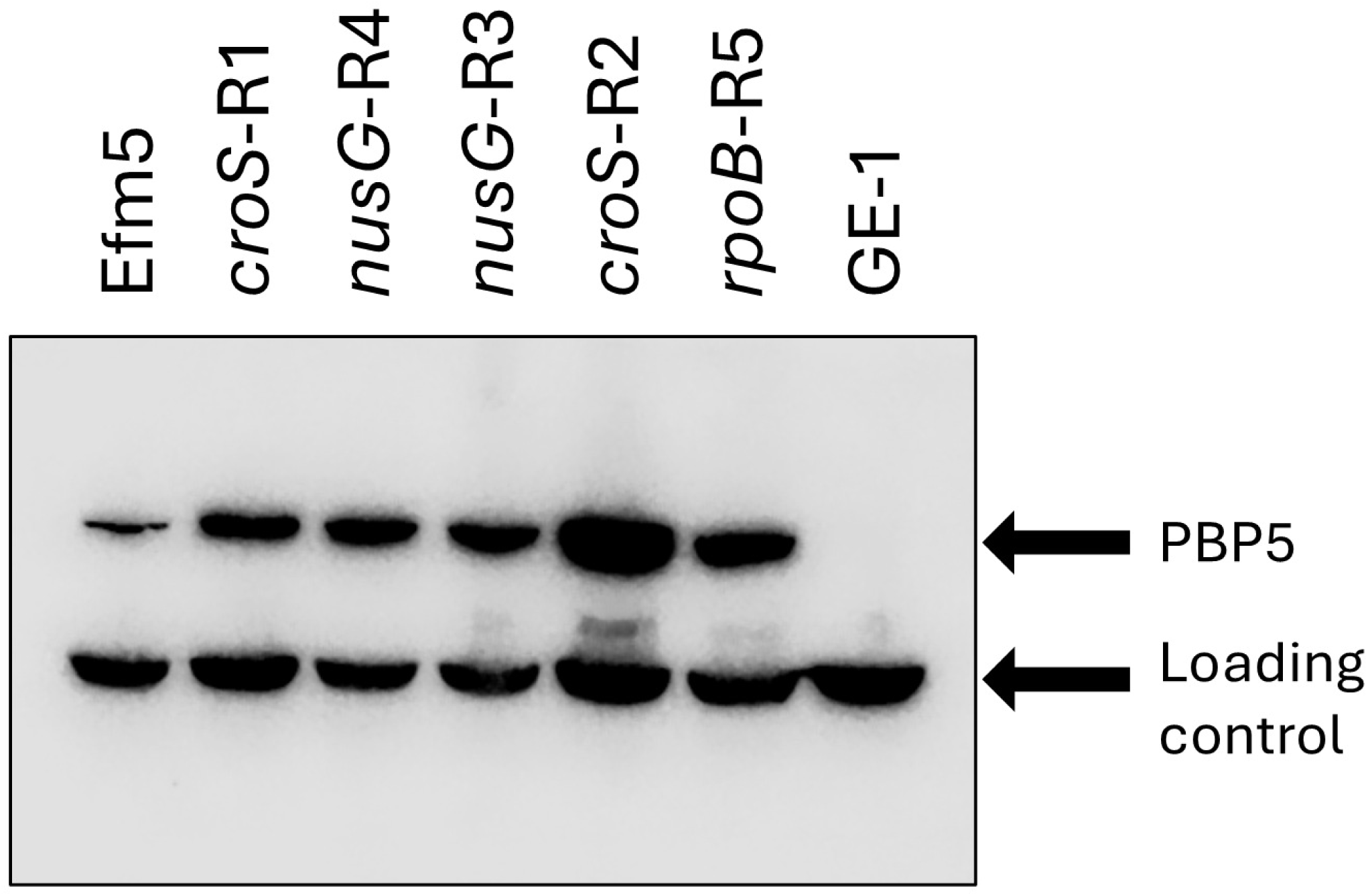
Western blotting of *E. faecium* Efm5, five CPH high-MIC variants and GE-1 (a *pbp5* negative mutant) using rPBP5-S polyclonal rat sera. In addition to the band corresponding to PBP5 (∼70 kDa), a non-specific lower-molecular-mass band was detected and used as an internal loading control.

The increased expression of PBP5 observed prompted us to investigate transcriptional changes in the CPH high-MIC variants. To this end, transcription of the entire *pbp5* operon was assessed by RT-qPCR. Our results revealed significant activation of the *ftsW*, *psr* and *pbp5* genes in the *croS-*R1 and R2 CPH high-MIC variants (Figure 4A). Transcription of *rutF*, located immediately upstream of the operon, remained unchanged. Additionally, no significant changes were observed in the transcript levels of *pbpA* or *croR*, although *croS* expression was slightly activated in the *croS-*R1 variant. In contrast, no changes in the *pbp5* operon were detected in the *nusG-*R3, *nusG-*R4 and *rpoB-*R5 variants, although these variants exhibited downregulation of *croS* and *croR*. The *nusG-*R4 and *rpoB-*R5 variants also showed slightly repressed expression of *ftsW*.

**Figure 4:**
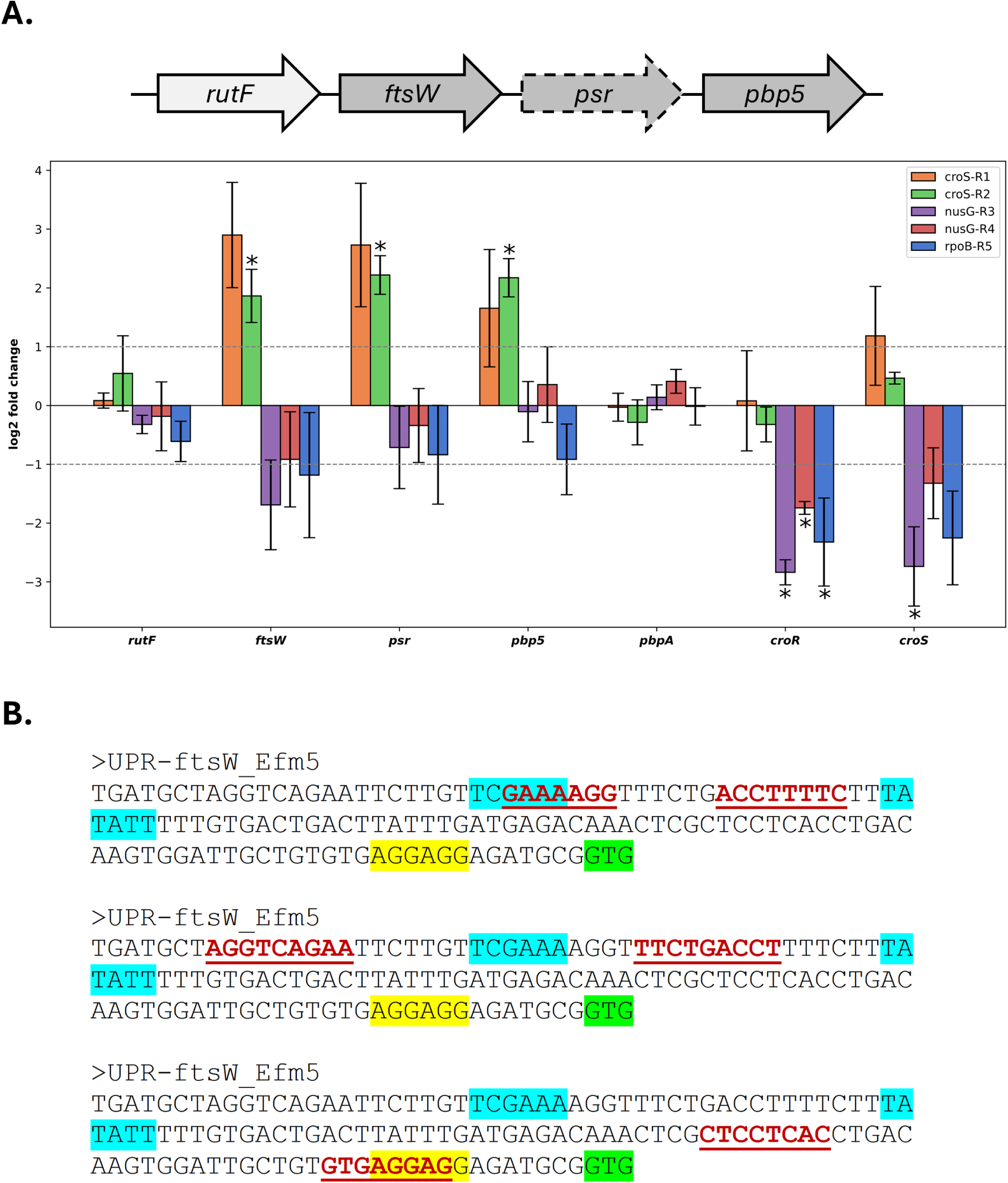
(A) Transcriptional analysis of the *pbp5* operon in CPH high-MIC variants compared to the *E. faecium* Efm5 isolate. Fold-change values (log₂) for the three genes of the *pbp5* operon (*ftsW, psr*, and *pbp5*), *rutF* (located immediately upstream of the operon), and the regulatory genes *croR* and *croS* are shown. Dashed lines indicate biologically relevant fold-change thresholds. Asterisks highlight genes with statistically significant expression differences relative to the Efm5 strain. (B) Schematic representation of the upstream region of the *ftsW* gene in Efm5, illustrating putative regulatory elements. The −35 and −10 promoter sequences are highlighted in light blue, the ribosome binding site (RBS) in yellow, and the translation start codon in light green. UPR, upstream region.

These findings prompted us to examine the upstream region of *ftsW*, the first gene of the *pbp5* operon, for potential regulatory sequences. The predicted −35 (TCGAAA) and −10 (TATATT) regions were located 101 and 76 nucleotides upstream of the *ftsW* translational start codon, respectively (Figure 4B). As it is typical for canonical promoters controlled by the vegetative sigma factor, the - 35 and −10 regions were spaced 19 bp apart. Additionally, several putative regulatory sequences, including direct and inverted repeats, were identified within the *ftsW* upstream region. Some of these sequences were situated near the −35 and −10 elements, suggesting a potential role in modulating RNA polymerase binding and, consequently, transcription of the *pbp5* operon. Furthermore, some regulatory sequences were found near or overlapping the ribosome binding site (RBS), which may indicate a role of these elements in post-transcriptional regulation.

### Analysis of cell morphology and ultrastructure

The morphology of Efm5 and a representative strain of each CPH high-MIC variant were examined by SEM (Figure 5A). As expected, all four strains exhibited pairs of cells (diplococci). Cocci chains were evident in the *nusG-*R3 and *rpoB-*R5 variants. No cell elongation was observed for Efm5 and CPH high-MIC variants, a feature typically associated with a shortage of peptidoglycan precursors (26).

**Figure 5:**
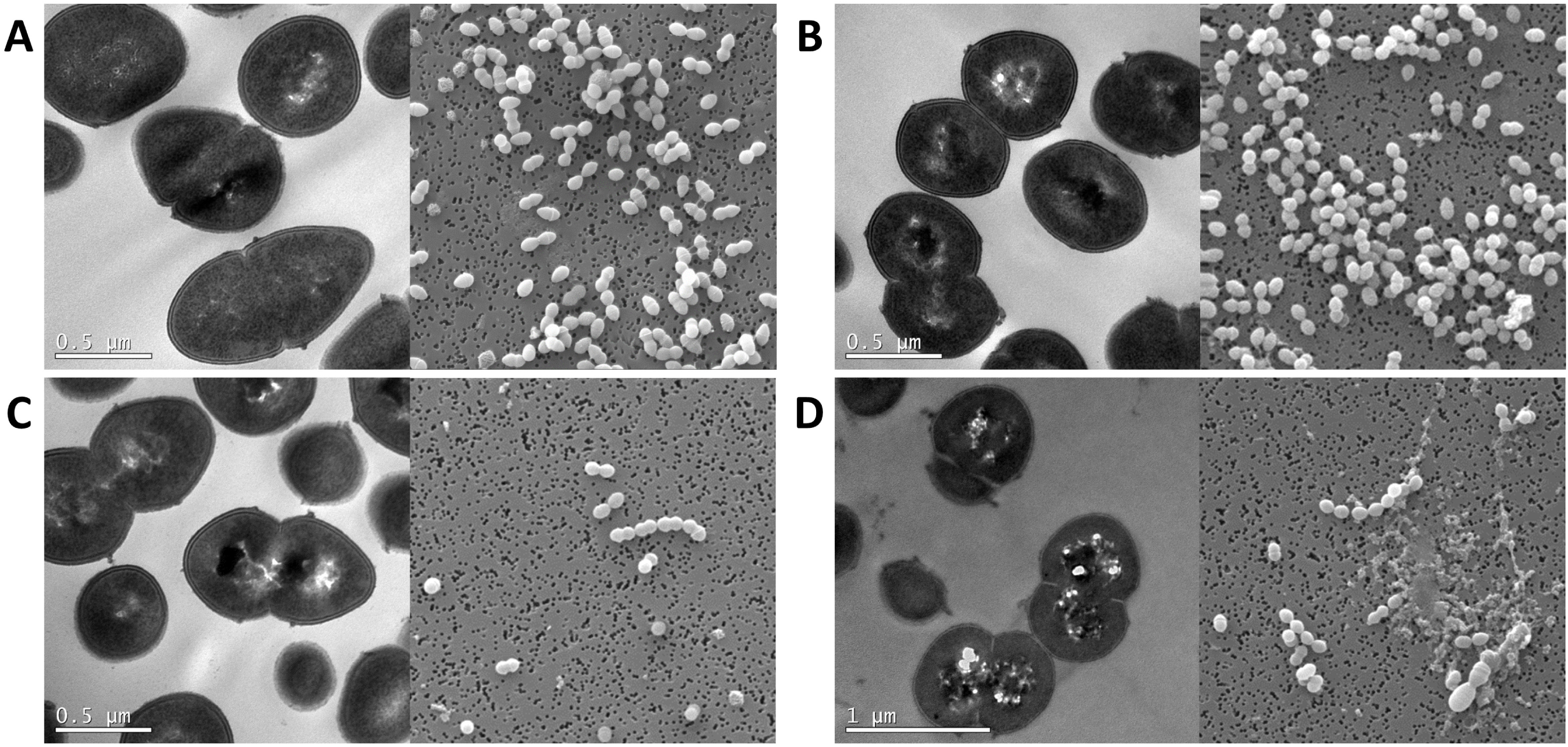
Scanning Electron Microscopy (SEM) and Transmission Electron Microscopy analyses of (A) Efm5, (B) *croS*-R1, (C) *nusG*-R3 and (D) *rpoB*-R5. All SEM figures are shown in the same magnification.

TEM was also performed to assess the ultrastructural features of the strains (Figure 5B). No significant differences in cell wall thickness were noted in Efm5 and CPH high-MIC variants. Unlike Efm5, all three variants displayed small round bodies with an electron-dense crossing structure, which was most prominent in the *nusG-*R3 variant. Additionally, the *rpoB-*R5 variant displayed electron-lucent cytoplasmic regions, appearing as clear zones. These regions were observed in independent preparations of this strain, further distinguishing it from the other CPH high-MIC variants.

## DISCUSSION

This study provides new insights into the genetic mechanisms underlying CPH resistance in *E. faecium*. Through comparative genomic analysis of the clinical isolate Efm5 (ST1195), which exhibited low CPH MICs, and several high-MIC derivatives selected under CTX pressure, we identified key mutations associated with CPH resistance and upregulation of cell wall-related genes. The CPH-susceptible phenotype of Efm5 challenges current assumptions about the intrinsic nature of CPH resistance in *E. faecium* and opens new perspectives on how this species may adapt to β-lactams in clinical settings.

By selecting high-level CPH-resistant variants from Efm5 under antibiotic pressure, we demonstrated that significant MIC increases can readily emerge through discrete genetic events. Notably, these shifts occurred without concomitant increases in ampicillin or ceftaroline resistance, pointing to alternative or additive resistance mechanisms beyond canonical alterations in PBP5. The identification of non-synonymous mutations in *croS*, *nusG* and *rpoB* across independent resistant variants underscores the multifactorial nature of CPH resistance in *E. faecium*.

In particular, the *croS* mutations V171A (*croS*-R1) and R343H (*croS*-R2) were located near the key phosphoryl-accepting residue H173, suggesting an impact on signal transduction within the CroRS two-component system, a known regulator of genes involved in lipid II biosynthesis, including the *pbp5* operon (6, 7). Consistent with a regulatory disruption, both CroS variants showed increased transcription of *pbp5*, *ftsW* and *psr*, along with elevated PBP5 protein levels, even in the absence of antibiotic pressure, suggesting a constitutive activation of the CroRS pathway driving cell wall remodeling strategies. The involvement of the CroRS system in CPH resistance has been previously reported in *E. faecalis* (5). Of note, recent studies have shown that the CroS extracellular domain is essential for proper signaling, yet it is not involved in ligand binding or direct sensing (27).

Unlike the CroS variants, the *nusG* and *rpoB* mutations appear to drive CPH resistance through distinct, possibly post-transcriptional mechanisms. These variants do not activate transcription of the *pbp5* operon and show a slight downregulation of *croS* and *croR*, along with minor repression of *ftsW*, suggesting an indirect regulatory effect. Nevertheless, they exhibit increased levels of PBP5, indicating an uncoupling between transcript abundance and protein expression. This pattern suggests that broader regulatory processes beyond the canonical *pbp5*-driven model may be involved in modulating resistance. Consistent with this, previous studies have shown that *rpoB* mutations (H486Y, H486D, and Q473K) can alter CPH susceptibility in *E. faecium* and *E. faecalis* (28), although the molecular basis remains unclear. The role of *rpoB* in this context aligns with earlier reports showing that specific mutations in this gene can influence CPH susceptibility in *Enterococcus* spp., even though the molecular basis remains unclear (28). Our findings add to this evidence and reinforce the notion that RpoB mutations contribute to antibiotic resistance beyond their well-known association with rifampicin, as also reflected in the observed increase in rifampicin MIC. Furthermore, recent studies have linked RpoB alterations to daptomycin resistance (29), underscoring the pleiotropic effects of this global transcriptional regulator.

The involvement of NusG is particularly intriguing, as this gene encodes a conserved transcription termination factor not previously implicated in antibiotic resistance. Its association with altered PBP5 expression introduces a novel link between transcriptional regulation and β-lactam susceptibility, warranting further investigation. The complexity of the *pbp5* promoter region, containing both canonical motifs and additional regulatory elements near the ribosome binding site, supports the existence of multilayered control mechanisms potentially sensitive to global regulatory perturbations.

Structural and morphological defects observed in all variants by SEM and TEM, most notably in the *nusG*-R3 and *rpoB*-R5 strains, which exhibited extensive chaining and electron-dense bodies, may reflect underlying disruptions in cell division or peptidoglycan remodeling. Such perturbations are consistent with stress-induced morphogenetic responses documented in enterococci and staphylococci (30, 31), implying that resistance-associated mutations could exert pleiotropic effects impacting cell envelope integrity. The presence of electron-lucent cytoplasmic regions specifically in the *rpoB*-R5 variant further suggests metabolic dysregulation or altered transcriptional activity, reinforcing the concept that these mutations induce complex physiological shifts beyond mere target modification.

Further studies are needed to elucidate the genetic basis of the low CPH MIC phenotype in *E. faecium*. To date, our group has identified four *E. faecium* isolates with this uncommon profile (Efm1, Efm5, Efm10 and Efm54) all obtained from patients with bacteremia (10). As described earlier, we found a premature stop codon in *psr* in the Efm5 isolate, raising the possibility that disruption of this gene may contribute to CPH susceptibility. Nonsense mutations in *psr* were also found in Efm1, which shares the low-MIC phenotype, and in Efm57, which instead displayed the classical intrinsic resistance to these antibiotics (19). The presence of low-MIC isolates without the premature stop codon (Efm10 and Efm54) further indicates that *psr* disruption alone is insufficient to explain the phenotype. Recently, it has been shown that clade A *E. faecium* isolates feature truncated *psr* alleles (757 bp vs. the typical 885 bp), which has been associated with high ampicillin MICs. However, Efm5 and the other three low-MIC isolates from our collection carried a full-length *psr* gene (885 bp), which is associated with low PBP5 expression. Therefore, although we cannot rule out a potential influence of *psr* variants in CPH susceptibility, our results suggest that, at least, additional genetic determinants are involved.

In conclusion, we identified previously uncharacterized mutations in *croS*, *nusG* and *rpoB* that demonstrate how CPH resistance in *E. faecium* can reemerge via multiple evolutionary paths, including modifications in signal transduction systems and global transcriptional regulators. Moreover, we demonstrate the existence of naturally occurring *E. faecium* with low-MICs toCPH, which will probably be considered as susceptible if breakpoints were available. The presence of these low-MICs isolates is interesting, and the genetic causes require further attention. Our findings carry important implications for surveillance and antibiotic stewardship, especially in clinical contexts where CPH are employed as part of combination therapy against multidrug-resistant enterococcal infections. Overall, our results advance current understanding of enterococcal adaptation to β-lactams and highlight the need to consider atypical resistance phenotypes in diagnostic and therapeutic strategies.

## ACKNOWLEDGEMENTS

This work was supported by grant PI24/01294 from Instituto de Salud Carlos III and co-founded by the European Union. The anti-PBP5 monoclonal antibody was a kind gift of Dr. Kavindra V. Singh (Division of Infectious Diseases, Department of Medicine, Houston Methodist Research Institute, Houston, Texas, USA). We also want to acknowledge the CERCA Program/Generalitat de Catalunya as well as FCT—Fundação para a Ciência e a Tecnologia, I.P., in the scope of the projects UIDP/04378/2020 and UIDB/04378/2020 of the Research Unit on Applied Molecular Biosciences— UCIBIO and the project LA/P/0140/2020 of the Associate Laboratory Institute for Health and Bioeconomy—i4HB.

---

**Table S1:** Primers used in this study.

**Table S2:** Homologs of CroS, NusG, and RpoB identified across 767 complete *Enterococcus* assemblies from NCBI using reciprocal BLASTP. The table includes protein identifiers and corresponding amino acid sequences.

